# Deep Prediction of Human Essential Genes using Weighted Protein-Protein Interaction Networks

**DOI:** 10.1101/2024.10.09.616990

**Authors:** Soroush Mehrpou, Eghbal G. Mansoori

## Abstract

Essential proteins are group of proteins that are indispensable to survival and development of cells. Prediction and analysis of essential genes/proteins are crucial for uncovering the mechanisms of cells. Using bioinformatics and high-throughput technologies, forecasting essential genes/proteins by protein–protein interaction (PPI) networks have become more efficient than traditional approaches which use expensive and time-consuming experimental methods. Previous studies have found that the essentiality of genes closely relates to their properties in PPI network. In this work, we propose a supervised deep model for predicting human essential genes using neighboring details of genes/proteins in the PPI network. Our approach implements a weight-biased random walk on PPI network to get the node network context. Then, some different measures are used to get some feature vectors for each node (gene/protein) that preserve the network structure as well as the gene’s properties in the PPI network. These feature vectors are then fed to a Relational AutoEncoder to embed the genes’ features into latent space. At last, these embedded features are put into a trained classifier to predict the human essential genes. The prediction results on two human PPI networks show that our model achieves better performance than those that only refer to genes’ centrality properties in the network.

## 1. Introduction

Essential genes are vital for the survival of a cell. Genes or their protein products interact with each other to perform various functions within the cell. Their interactions in the protein-protein interaction (PPI) network play a crucial role in predicting essential genes. Identifying essential genes in eukaryote and prokaryote organisms has contributed to understanding fundamental cellular requirements [1]. Additionally, in the field of synthetic biology, it has paved the way for research on constructing cells with minimal genomes. Previous studies indicate a close relationship between essential genes and human diseases. Therefore, the identification of essential genes has a significant role in understanding the minimal requirements for cells’ survival and discovering medicine for diseases [2].

Up to date, an extensive works has been done to predict essential genes using experimental biological methods and network-based approaches. Although experimental laboratory methods such as gene knockout [3], RNA interference [4], and conditional gene knockout [5] could provide reasonably accurate predictions of essential genes, they were time-consuming and costly. Nowadays, with the advancement of high-throughput technologies such as yeast two-hybrid systems [6] and mass spectrometry-based proteomics [7], various PPI data are available which are used in the study of essential genes.

The research results demonstrate a correlation between the topological features of PPI networks and the essentiality of genes/proteins. It has also been revealed that in some cases, proteins that strongly interact with other proteins in the network are more likely to be important compared to proteins with fewer connections [8]. Therefore, centrality measures are widely used in the analysis of PPI networks to assess the importance or centrality of nodes (genes) in the network. Since most essential genes/proteins are conservative, some methods have been suggested that combine protein ortholog information, such as SON [9]. Several studies detected essential genes/proteins based on weighted PPI networks [10]. Some researchers have made significant efforts to improve the essential gene prediction accuracy in some simple eukaryotic organisms by employing machine-learning methods and combining some topological and biological features.

As mentioned above, the protein interactions in PPI networks play a crucial role in predicting essential genes. Recently, with the advancement of biological techniques, PPI networks have become more comprehensive and accurate. As for human essential gene identification, we can draw a lesson from previous studies on other species. For this purpose, we try to investigate the importance of genes in graph nodes by analyzing the neighbors of each node at higher levels to obtain a more global view of the graph. So, proposing an effective method to extract important features of genes in PPI networks is our first goal.

In a biased random walk through a graph, the probability of moving from one node to another is not uniform but biased in some ways [11]. As our second goal in this work, we get more neighbors’ level of each node (gene) in human weighted PPI networks by using a weight-biased random walk method which walks more likely to those connecting nodes sharing similar neighbors. To build a feature vector for each protein, we use some topological and biological measures of neighboring proteins in the PPI networks. Then, these features are fed into a Relational AutoEncoder as a new network embedding to map the genes’ features into latent space while maximally preserving the relationship between the genes and their neighbors in the PPI networks. At last, the genes’ features are put into a trained classifier to predict the essential genes.

The rest of this paper is arranged as follows. The related work on essential genes prediction is reviewed in Section 2. Section 3 explains our proposed method. Section 4 presents the experiments on our method and the compared ones. The paper is concluded in Section 5.

## 2. Related Works

As mentioned before, genes tend to be essential due to their high connectivity and central position in the PPI network. Some centrality methods are frequently used to measure the importance of the genes in the PPI network, such as degree centrality (DC) [12], closeness centrality (CC) [13], betweenness centrality (BC) [14], information centrality (IC) [15], eigenvector centrality (EC) [16], subgraph centrality (SC) [17] and edge clustering coefficient (NC) [18].

Li et al. and Peng et al. proposed two methods for identifying essential genes/proteins by combining PPI networks and gene expression data, namely PeC [19] and WDC [20], respectively. Some studies indicate that essential genes/proteins tend to gather more in protein complexes [21].

Based on this perspective, two modified methods called UC and UC-P [22], which integrate information from protein complexes, were proposed by Li et al. for identifying essential genes/proteins. Additionally, recent studies suggest that subcellular localization of proteins may play a crucial role in identification of essential genes/proteins. Tang et al. proposed a method called CNC [23], which integrates subcellular localization information to enhance the accuracy of essential gene/protein identification.

In some studies, essential genes/proteins are detected based on weighted PPI networks. Xu et al. proposed a method named essentiality ranking [10] that integrates multiple data sources to weighted PPI networks. Recently, Peng et al. proposed a new prediction method, named UDoNC [24], by combining the domain features of proteins with their topological properties in PPI networks. Lei et al. presented a new computational method with HITS algorithm on weighted PPI networks to identify essential genes/proteins, named HSEP [25].

Some machine-learning methods combine the topological and biological features to improve the essential gene prediction accuracy. For example, Chen et al. [26] trained an SVM classifier by the features of some known yeast essential genes/proteins, including the combination of phyletic retention, proteins’ evolutionary rate, paralogs, protein size, and centrality degree in PPI networks and gene-expression networks. Then, the classifier was adopted to predict other potential yeast essential genes. Zhong et al. [27] developed an XGBoost-based classifier to integrate multiple topology properties of essential genes in the PPI network to predict essential genes of yeast and E. coli.

Zhang et al. [28] proposed a new method by constructing a dynamic PPI network by identifying active genes at each time point from gene expression profiles. They designed a novel centrality method to assign each gene a ranking score in the final network while considering its orthologous property and its global/local topological properties in the network.

Dai et al. [11] proposed a word2vec embedding method on PPI network. For prediction, they trained a machine learning model on genes’ feature vectors. They also implemented a bias random walk on the network to get the node network context. In another research, He et al. [29] developed a new essential gene detection method which extracts features from weighted protein-domain network. This network is constructed by combining known protein-protein interactions with known associations between proteins and domains.

## 3. The Proposed Model

Our human essential gene identification model consists of some phases. It first uses biological information of gene ontology and protein sequences to weight the edges of genes/proteins in the PPI network. We introduce a novel weight-biased random walk algorithm which walks, randomly according to the weighted-edges, in the PPI network and visits the genes sharing similar neighbors. Using the visited genes, our approach extracts some topological and biological features of neighboring proteins in the PPI network and feeds them into a Relational AutoEncoder to embed into latent space while preserving the neighbors’ relationships. At the last stage, it trains a classifier to predict the essential genes. This classifier is designed in two scenarios (under-sampling and over-sampling) to handle the imbalanced data of human essential genes. Algorithm 1 gives the steps of our human essential gene identification model in pseudocode.

### Algorithm 1: Proposed model for identification of human essential genes

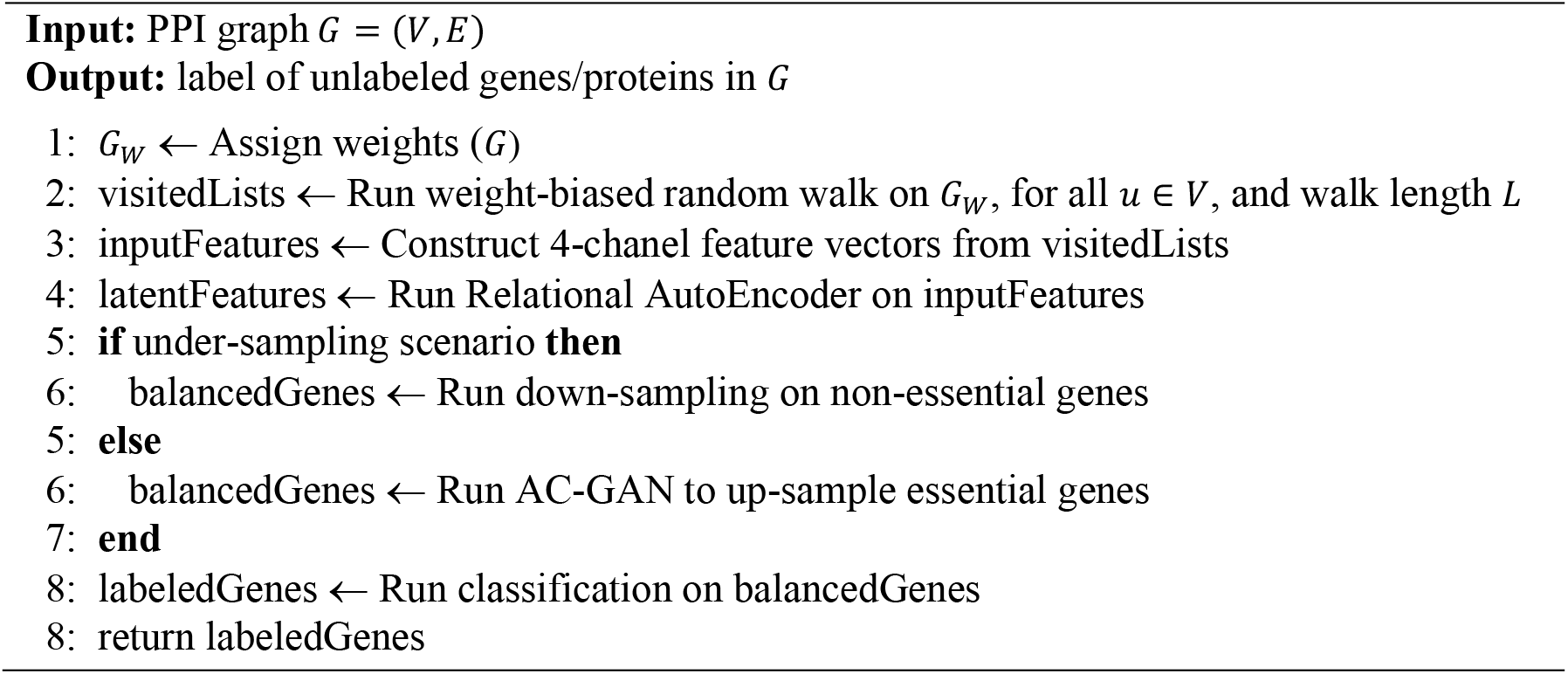

Figure 1 shows an overview of our model. The details of each stage are explained in the following subsections.

**Figure 1.**
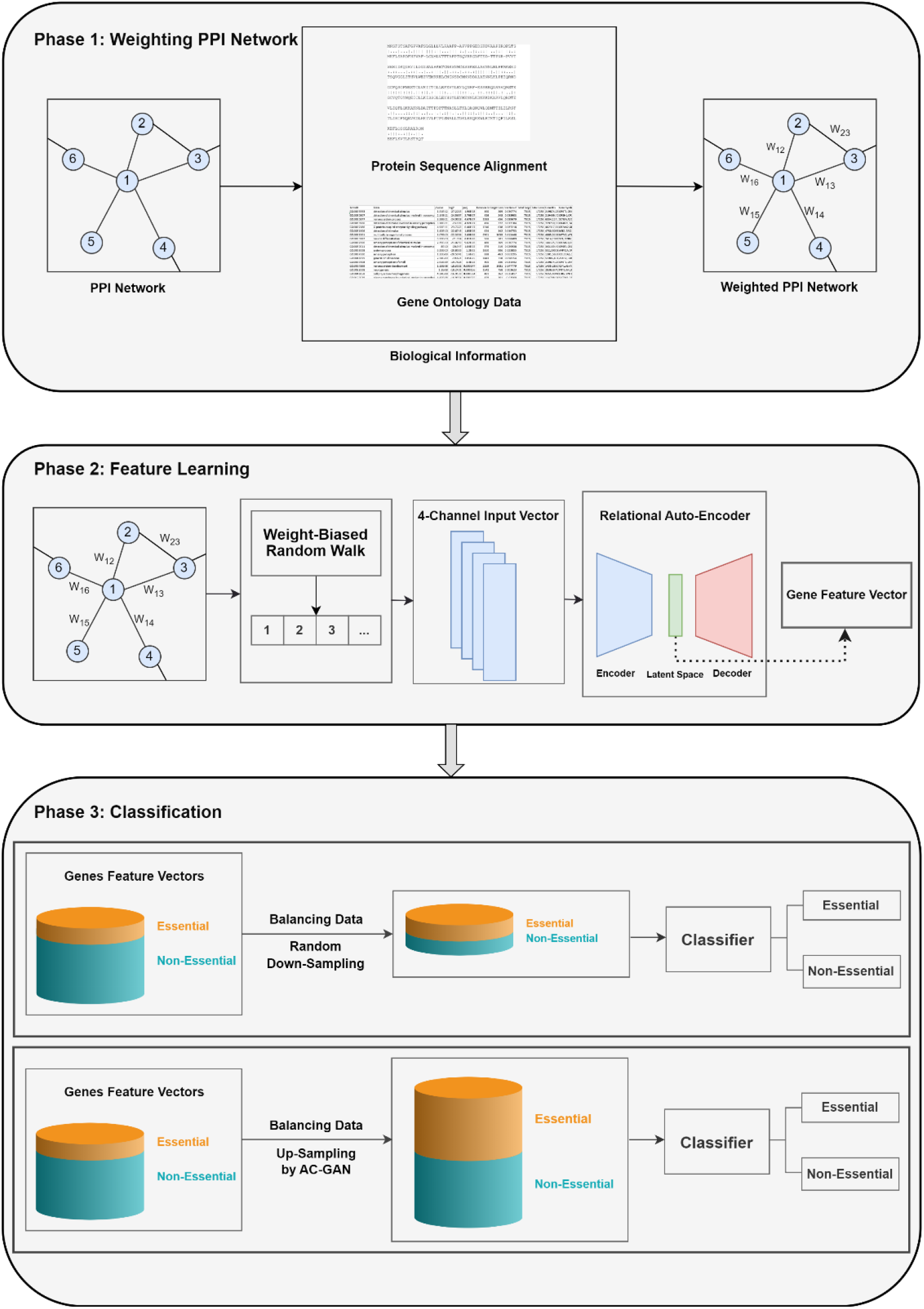
An overview of our proposed model for identifying human essential genes.

### 3.1. Constructing Weighted Protein-Protein Interaction Network

In order to show the reliability of connecting links in the PPI networks, we try to assign suitable weights to the links in two different schemes. In the first scheme to set the weight of link between two proteins, we pairwise align their sequences via lining up to achieve maximal levels of identity (and conservation) or similarity degree and homology possibility.

In the second schemes, the weights are assigned from prospective of proteins function similarity. In this regard, the gene semantic similarity between gene ontology (GO) terms of two proteins is calculated and is set as their link score. Indeed, if proteins have similar annotations, the similarity score increases and the connection is stronger. GO annotation is a statement that created by associating a gene or gene product with GO terms. To evaluate the similarity of proteins according to proteins GO terms in the PPI network, we used the method in [30]. The GO semantic similarity between proteins *A*and *B*is computed as:

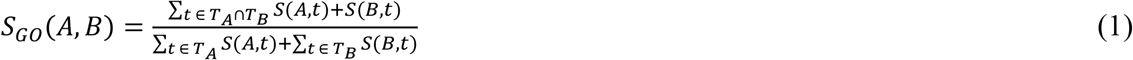

where *T*_*A*_denotes the GO terms of protein *A*, and *S*(*A, t*) computes the S-value of GO term *t* in *T*_*A*_.

In this paper, we use both schemes in some ways according to the length of two protein sequences, connected in PPI network. If these sequences are close in length (their length difference is less than a prespecified threshold), their pairwise alignment score is used for the weight of their link. Otherwise, we adopt their GO semantic similarity in (1) to calculate their weight.

### 3.2. Weight-Biased Random Walk

A biased random walk is a type of random walk in a graph where the probability of moving from one node to another is not uniform but biased in some ways. In other words, it’s a stochastic process where you start at a node in a graph and, at each step, choose a neighbor to move to, but the choice is influenced by certain biases or preferences [11]. In this way, by setting a length for random walk algorithm, it is possible to analyze the neighbors in more levels of distance on the graph.

Weighted random walk will manage the nodes’ selection based on their connection strength. While candidate connection with a higher weight score will be less likely to be eliminated, there is still a chance that they may be eliminated because their probability of selection is less than 1. In this work, we introduce a novel weight-biased random walk which walks more likely to those connecting nodes that share similar neighbors. Using it, we get more neighbors’ level of each node in human weighted PPI networks. In order to get features of each node based on its neighbors, it is possible to analyze the neighbors in further levels.

We denote a weighted PPI graph as *G*_*W*_ = (*V, E, W*) where *V* and *E* denote the set of vertices and edges respectively, and *W* is the weight set of edges, showing reliability of connections. The probability of walking from node *v* ∈ *V* to another node *t* ∈ *V* is calculated, based on cumulative probability distribution, using weights of *v*’s neighbors:

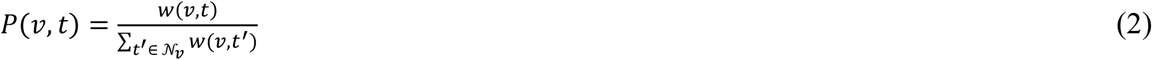

where 𝒩_*v*_ is the neighbors of node *v*.

For walking from current node (gene/protein) *v* to the next node in its neighbors (i.e., 𝒩_*v*_), we use the Roulette Wheel selection mechanism [31]. It selects the next node *u* ∈ 𝒩_*v*_, randomly according to the weights of its neighboring nodes and walks to this new node (gene/protein). Starting from seed node *u*, this process repeats until *L* walks is done, where *L* is the walk length. Algorithm 2 gives the pseudocode of our weight-biased random walk.

#### Algorithm 2: Weight-biased random walk algorithm

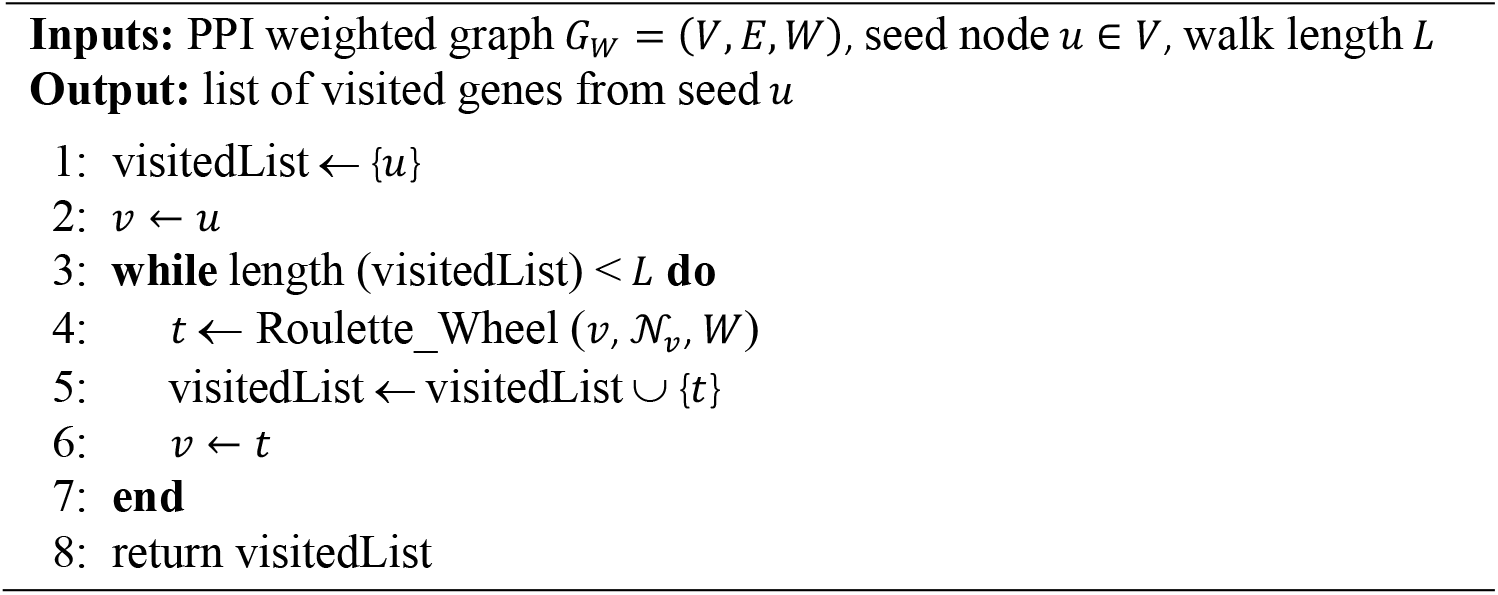

### 3.3. Feature Learning

After getting the list of visited neighbors for each protein in the PPI network (via running the weight-biased random walk algorithm), we construct a 4-channel input feature space from these lists for each PPI node. These features are then fed to a deep Relational AutoEncoder to embed the feature vector of proteins into the latent space. This AutoEncoder uses 1D convolution layer to capture the features of each PPI node.

To construct the input feature space of the first channel, we measure the centrality of each node according to its neighbors. This centrality degree of proteins has been widely used as a good measure for predicting essentiality, in the literature [12] as named degree of centrality (DC). Since our PPI networks are now weighted, we use weighted degree of centrality (WDC) instead. The WDC for node *v* computes the total weights of the edges connecting node *v* to its neighbors in 𝒩_*v*_. That is:

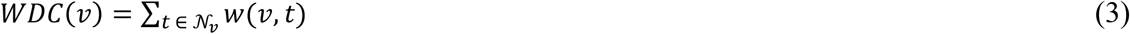

For the second channel, we calculate the shortest distance between each node (as seed) and its visited neighbors in the PPI network. In this way, we measure the farness of walk-outs from the seed nodes, which in turn reveals the level of neighborhood.

The third channel of input feature space reflects the essentiality of PPI nodes according to the labels of training data. From this perspective of proteins’ interactions in the functional modules, whether or not there are essential genes/proteins in the neighborhood of a protein, this can affect the essentiality of that protein. However, there are a few proteins in PPI network which have essential/non-essential labels, leaving the most unlabeled. For each seed node, we examine the labels of its visited neighbors in the PPI network and use to form the third channel. In this regard, the essential, non-essential and unknown nodes are encoded by 1, -1, and 0, respectively.

At last, we use ID of visited neighbors of each gene in the PPI network to form the fourth channel. In this way, it helps the genes with similar neighbors to have close feature vectors.

### 3.4. Relational AutoEncoder

In traditional autoencoders, the encoder module often does feature extraction via nonlinear transforming data from the original (often high dimensional) space to a latent (relatively lowdimensional) space. Our proposed Relational AutoEncoder also transforms nonlinearly the input feature vector into a code vector with lower dimensions, while minimizing the reconstruction loss of both data and their relationships. Its objective function can be defined as:

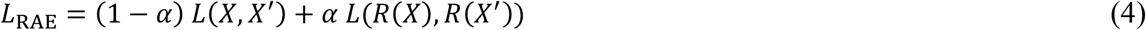

where *X* and *X*^’^ represent the training data and those reconstructed by AutoEncoder, respectively. Also, *R*(*X*) computes the relationship among data samples in *X*. In [32], data relationship is modelled by their similarities, obtained via multiplication of *X* and *X*^*T*^. The scale parameter *α* controls the weights of reconstruction loss and relationship reconstruction loss. Supposing Θ as neural network parameters of the autoencoder, the optimum Θ_opt_ is obtained via minimizing the loss function in (4) as [32]:

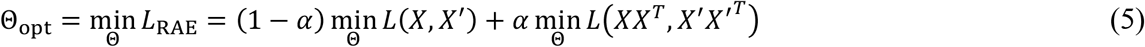

where loss function *L*(·, ·) uses the squared error.

The aim of using Relational AutoEncoder is that the nodes (genes) close in the PPI network would have similar feature representations in the latent space. Figure 2 shows the structure of our Relational AutoEncoder which consists of ConvNet, flatten, reshape, and DeConNet layers.

**Figure 2.**
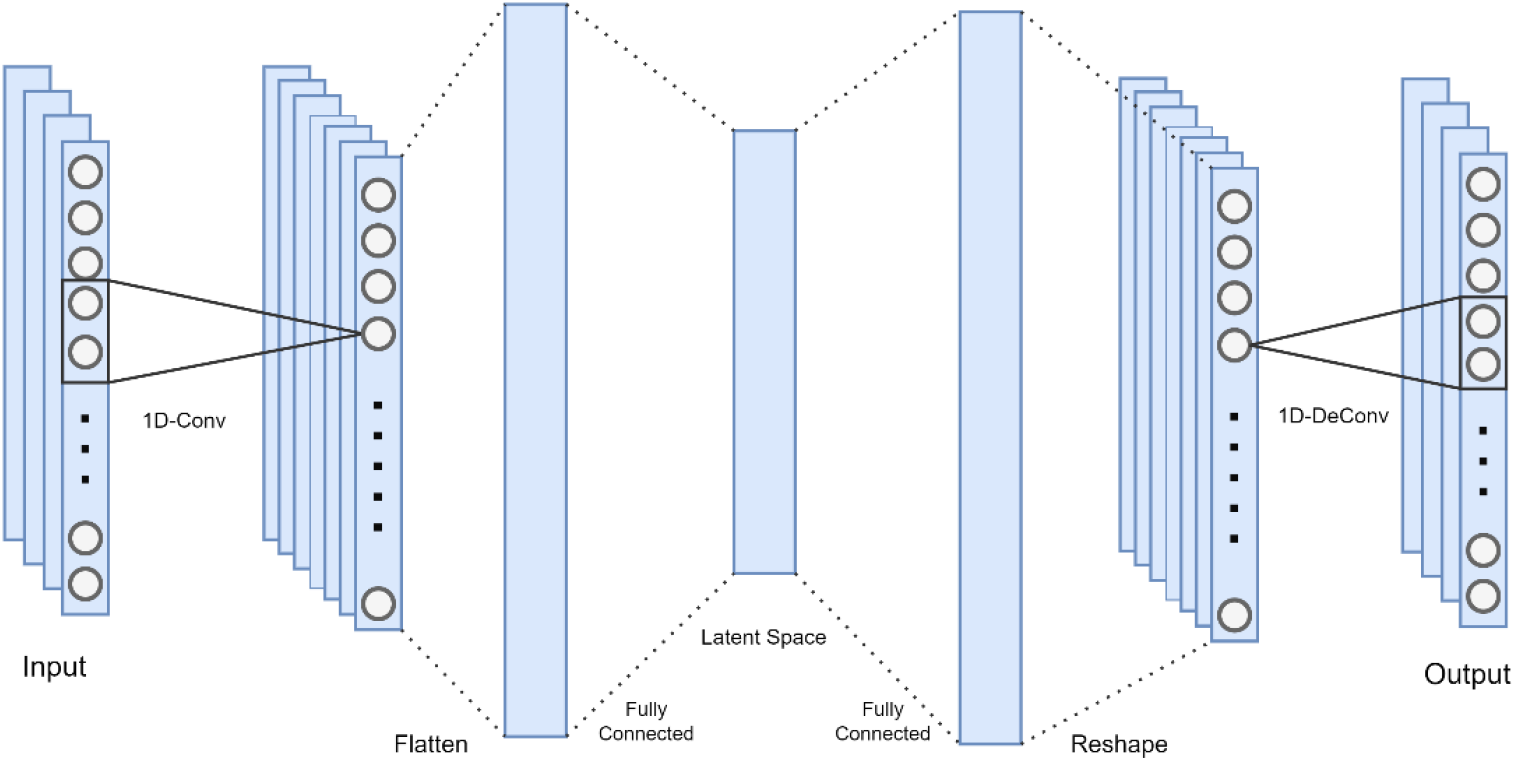
The structure of our Relational AutoEncoder.

### 3.5. Balancing Labeled Genes

As mentioned before, since the essential genes in the training data are imbalanced, we design the classifier, in Figure 1, for two distinct scenarios of under-sampling and over-sampling. For the under-sampling strategy, by supposing uniform distribution for non-essential data samples, they are down-sampled to keep 1:1 ratio for essential and non-essential gene/protein sequences.

On the other hand, for over-sampling scenario, we use an extension of Generative Adversarial Network (GAN), called Auxiliary Classifier GAN (AC-GAN) model [33], to generate synthetic data samples. Since non-essential samples are more abundant in training data, AC-GAN generates new samples similar to essential ones to balance the samples of essential and non-essential genes/proteins. The objective function of AC-GAN has two parts [33]: the loglikelihood of the correct source, *L*_*S*_, and the log-likelihood of the correct class, *L*_*C*_. Since Generator of AC-GAN generates labeled samples, the parts of its objective function are defined as:

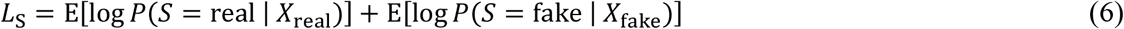

And

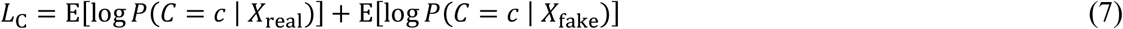

where probability distribution *P*(*S*|*X*) = *D*(*X*), and *D*(·) denotes the Discriminator function in GAN. Discriminator is trained to maximize *L*_*C*_ + *L*_*S*_ while Generator is trained to maximize *L*_*C*_ − *L*_*S*_.

### 3.6. Classification

In the last phase of the proposed model for essential gene/protein identification, as mentioned before, some classifiers are learned using the balanced training gene/protein sequences. These classifiers are then used to determine the essentiality/non-essentiality of the unlabeled genes/proteins. We aim to verify that the extracted/embedded feature vectors of genes in the PPI network is helpful for predicting the human essential genes. In this work, we used some popular classifiers for prediction task. These common classifiers include random forest [34], support vector machine with RBF kernel [26], decision tree [35], logistic regression [36], Naive Bayes [37], and *k*-nearest neighbor [38].

## 4 Experiments

In this section, we first introduce the datasets, used in the experiments. Then, the parameter setting of our algorithms is given. Also, the evaluation metrics, used in the experiments, are described. At last, the experimental results of our model against four state-of-the-art essential protein prediction algorithms are reported. Also, results of our model on different classifiers are discussed.

### 4.1. Datasets

In the experiments, we evaluated our model on two homo sapiens PPI networks. The FIs dataset was downloaded from Reactome database [39]. It has 12,277 genes/proteins and 230,243 interactions. The other dataset is InWeb-IM, downloaded from InBio Map database [40], with 17,428 genes/proteins and 625,641 interactions. Additionally, the supplementary dataset of human essential genes [41] is used for labelling, as essential/non-essential, the known genes/proteins of both datasets. This supplementary dataset consists of 1,516 essential and 10,499 non-essential genes.

Table 1 summarizes the characteristics of two FIs and InWeb-IM PPI networks, in addition to statistics of their labels.

**Table 1.** Statistics of PPI datasets.

| PPI network | # Genes | # Interactions | # Essential genes | # Non-essential genes |
| --- | --- | --- | --- | --- |
| FIs | 12,277 | 230,243 | 1,359 | 5,388 |
| InWeb-IM | 17,428 | 625,641 | 1,512 | 9,036 |

### 4.2. Experiment Setting

The walk length parameter, *L*, decides the walk distance in the PPI network. In order to find the data-driven walk length in our experiments, we run our weight-biased random walk algorithm on both FIs and InWeb-IM datasets for various walk lengths, and computed the area under curve of receiver operating characteristic (AUC-ROC) and area under curve of precision-recall (AUC-PR). The higher these areas are, the more reliable our algorithm is. Figure 3 shows these areas for both datasets. Accordingly, in the experiments, we set the walk length of weight-biased random walk algorithm to 10, for both datasets.

**Figure 3.**
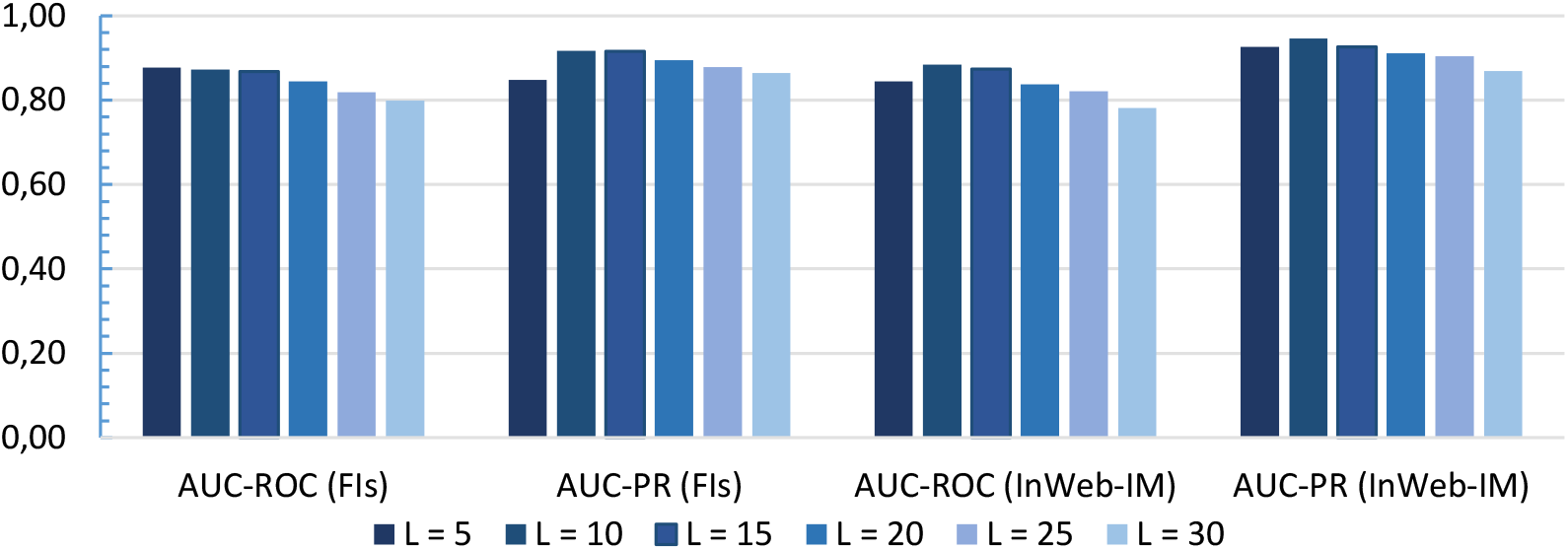
Effect of walk length on weight-biased random walk algorithm.

In our Relational AutoEncoder model, we preset the latent space size to 16, and the learning rate to 0.001, for both datasets.

In this study, we used 5-fold cross-validation to evaluate the prediction performance of our model and the competing methods. In this way, all genes in each dataset were randomly partitioned into five parts. Each time, one part set aside for test and the remaining genes were used for training the model. Repeating this steps five times, all parts were used up as test set, and the predicting results on test sets were averaged.

### 4.3. Evaluation Metrics

To further evaluate the performance of our proposed model, we used several statistical measures, namely sensitivity (*SN*), specificity (*SP*), positive predictive value (*PPV*), negative predictive value (*NPV*), F-measure, and accuracy (*ACC*), in the experiments. These criteria were also used for comparing our model against four state-of-the-art methods in essential protein prediction. These statistical measures are defined as follows.

In these criteria, *TP* (true positive) counts the number of truly predicted essential genes while *FP* (false positive) counts the number of non-essential genes that are predicted as essential. Similarly, *TN* (true negative) counts the number of truly predicted non-essential genes while *FN* (false negative) counts the number of essential genes that are predicted as non-essential. Also, *P* and *N* denote the number of essential and non-essential genes, respectively.

**Sensitivity**: the ratio of genes, accurately predicted as essential, to the total number of essential genes.

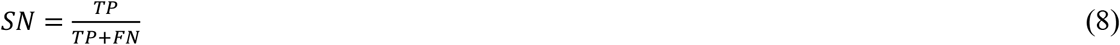

**Specificity**: the ratio of accurately predicted non-essential genes to the total number of non-essential genes.

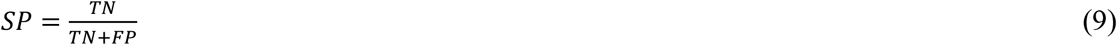

**Positive predictive value**: the ratio of genes accurately predicted as essential.

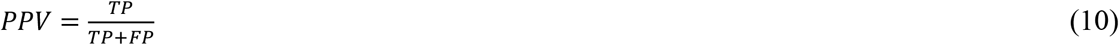

**Negative predictive value**: the ratio of genes accurately predicted as non-essential.

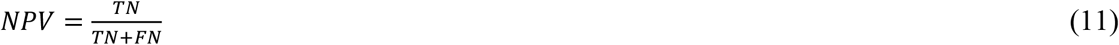

**F-measure**: the harmonic mean of *SN* and *PPV* is referred as F-measure.

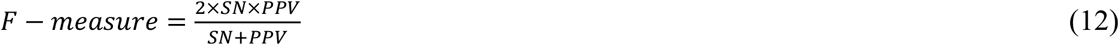

**Accuracy**: the ratio of correctly predicted genes, regarded as essential or non-essential, to all genes.

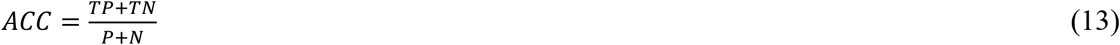

### 4.4. Classifier Selection

As mentioned in Algorithm 1 and depicted in Figure 1, the last phase of our model uses classifiers to predict the human essential genes. In order to select the suitable classifier, which achieves the best prediction using the learned features, we run six frequently-used classifiers, beside to Artificial Neural Network (ANN), in our model. These classifiers include *k*-nearest neighbor (KNN), Naive Bayes (NB), logistic regression (LR), support vector machine (SVM) with RBF kernel, decision tree (DT), and random forest (RF).

Table 2 compares the performance of our model, on FIs dataset, when uses different classifiers in its last phase. In this table, the results of both balancing methods, under-sampling and over-sampling, are included, where the best classifier in each evaluation metric is shown in boldface. According to the results, when random down-sampling of non-essential genes is applied to balance the FIs dataset, the RF classifier, as the last phase of our model, achieves the highest values in *SN, NPV, F* − *measure*, and *ACC* criteria, while for *SP* and *PPV* measures, the results of KNN are the best. On the other hand, the ANN classifier gives the best values when AC-GAN is used to up-sample the essential genes to balance this dataset.

**Table 2.** Performance of our model on FIs dataset with different classifiers.

| Classifier of<br>Our Model | Evaluation Metric |  |  |  |  |  |
| --- | --- | --- | --- | --- | --- | --- |
|  | SN | SP | PPV | NPV | F-measure | ACC |
| **Under-sampling** |  |  |  |  |  |  |
| ANN | 0.869 | 0.873 | 0.876 | 0.866 | 0.873 | 0.871 |
| KNN | 0.773 | **0.911** | **0.900** | 0.795 | 0.832 | 0.841 |
| NB | 0.888 | 0.827 | 0.841 | 0.878 | 0.864 | 0.858 |
| LR | 0.869 | 0.842 | 0.85 | 0.862 | 0.860 | 0.856 |
| SVM | 0.869 | 0.881 | 0.883 | 0.867 | 0.876 | 0.875 |
| DT | 0.869 | 0.854 | 0.860 | 0.864 | 0.865 | 0.862 |
| RF | **0.892** | 0.865 | 0.872 | **0.886** | **0.882** | **0.879** |
| **Over-sampling** |  |  |  |  |  |  |
| ANN | **0.812** | 0.941 | 0.825 | **0.937** | **0.818** | **0.909** |
| KNN | 0.655 | **0.960** | 0.849 | 0.892 | 0.739 | 0.883 |
| NB | 0.741 | 0.958 | **0.857** | 0.916 | 0.795 | 0.903 |
| LR | 0.756 | 0.953 | 0.845 | 0.920 | 0.798 | 0.903 |
| SVM | 0.782 | 0.941 | 0.819 | 0.927 | 0.800 | 0.901 |
| DT | 0.734 | 0.924 | 0.765 | 0.911 | 0.749 | 0.876 |
| RF | 0.786 | 0.934 | 0.801 | 0.928 | 0.793 | 0.897 |

The above experiments on FIs dataset were also repeated for InWeb-IM dataset. Table 3 shows their results. In this case, for both down-sampling and up-sampling approaches of balancing, the ANN classifier gives the best values in most of the criteria, though for some measures, KNN and SVM perform the best. Therefore, the ANN classifier was used as the last phase of our model, in the next experiments.

**Table 3.**
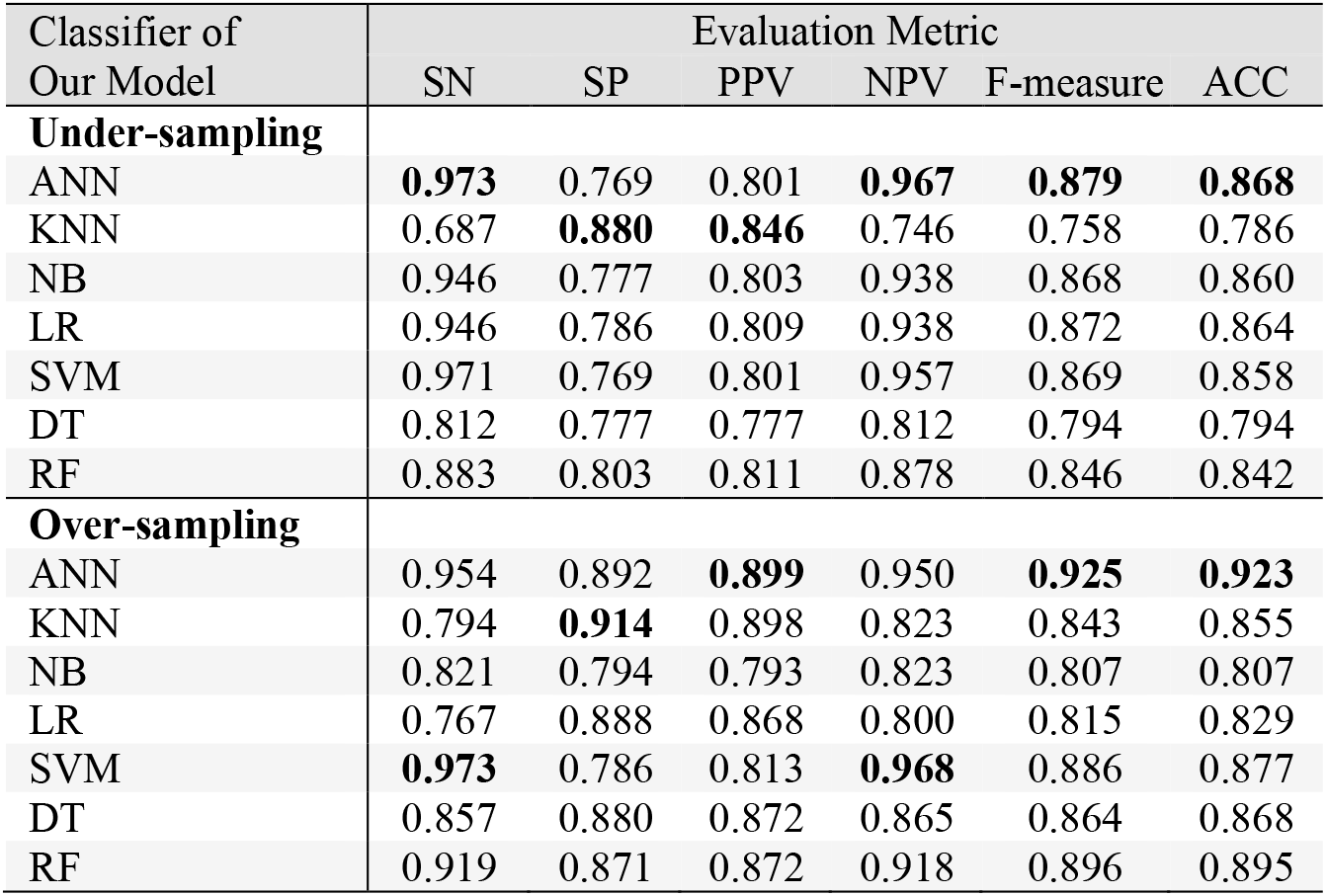
Performance of our model on InWeb-IM dataset with different classifiers.

### 4.5. Model Comparison

In order to assess the performance of our model against some state-of-the-art methods, we compared it with four existing approaches in the literature, including the centrality-based method, the Word2Vec method [11], the DeepWalk method [42], and the LINE method [43]. As mentioned before, the centrality-based method uses different centrality measures of genes in the PPI network as input features. These centrality measures include DC, BC, CC, SC, EC, IC, and NC. On the other hand, the Word2Vec, DeepWalk, and LINE methods, as well as our model, learn the feature representation for every gene in the human PPI network by different network embedding algorithms. Additionally, as concluded from the experimental results in the previous subsection, we used the ANN classifier, in all methods, to make predictions.

For under-sampling balancing method, we randomly selected 1,359 essential and 1,512 non-essential genes, for both FIs and InWeb-IM datasets. Table 4 shows the performance of our model against the compared methods, on both FIs and InWeb-IM datasets, where the best method in each evaluation metric is shown in boldface. Clearly, our model achieves the best results in most of the criteria, except for *SP* and *PPV* measures where Word2Vec and/or Centrality act better. This exception might be resolved if ANN classifier is substituted by KNN, according to the previous results.

**Table 4.** Performance of our model against the rival methods when the data balanced using random under-sampling.

| Method | Evaluation Metric |  |  |  |  |  |
| --- | --- | --- | --- | --- | --- | --- |
|  | SN | SP | PPV | NPV | F-measure | ACC |
| **FIs Dataset** |  |  |  |  |  |  |
| Centrality | 0.553 | **0.904** | 0.852 | 0.669 | 0.671 | 0.728 |
| LINE | 0.847 | 0.647 | 0.705 | 0.808 | 0.770 | 0.747 |
| DeepWalk | 0.846 | 0.830 | 0.833 | 0.843 | 0.839 | 0.838 |
| Word2Vec | 0.836 | 0.863 | 0.859 | 0.840 | 0.847 | 0.849 |
| Our Model | **0.869** | 0.873 | **0.876** | **0.866** | **0.873** | **0.871** |
| **InWeb-IM Dataset** |  |  |  |  |  |  |
| Centrality | 0.713 | 0.839 | 0.816 | 0.745 | 0.761 | 0.776 |
| LINE | 0.905 | 0.492 | 0.641 | 0.839 | 0.750 | 0.699 |
| DeepWalk | 0.848 | 0.811 | 0.817 | 0.842 | 0.832 | 0.829 |
| Word2Vec | 0.858 | **0.854** | **0.855** | 0.858 | 0.857 | 0.856 |
| Our Model | **0.973** | 0.769 | 0.801 | **0.967** | **0.879** | **0.868** |

After using AC-GAN to over-sample the essential genes of FIs dataset, we obtained the balanced data of 5,388 essential and 5,388 non-essential genes. For InWeb-IM dataset, the balanced dataset now contains 9,036 essential and 9,036 non-essential genes.

The performance of our model against the competing methods is compared in Table 5, where the best method in both datasets and each evaluation metric is shown in boldface. For both datasets, our model achieves the best results in all criteria, except *SP* where the LINE method performs better. This also occurs in *PPV* metric for FIs dataset. Again, using KNN instead of ANN, this preference might be changed.

**Table 5.** Performance of our model against the competing methods when AC-GAN was used to over-sample the data.

| Method | Evaluation Metric |  |  |  |  |  |
| --- | --- | --- | --- | --- | --- | --- |
|  | SN | SP | PPV | NPV | F-measure | ACC |
| **FIs Dataset** |  |  |  |  |  |  |
| Centrality | 0.523 | 0.917 | 0.608 | 0.879 | 0.562 | 0.836 |
| LINE | 0.394 | **0.981** | **0.837** | 0.866 | 0.539 | 0.856 |
| DeepWalk | 0.621 | 0.954 | 0.771 | 0.909 | 0.688 | 0.887 |
| Word2Vec | 0.597 | 0.968 | 0.822 | 0.905 | 0.692 | 0.893 |
| Our Model | **0.812** | 0.941 | 0.825 | **0.937** | **0.818** | **0.909** |
| **InWeb-IM Dataset** |  |  |  |  |  |  |
| Centrality | 0.562 | 0.916 | 0.527 | 0.926 | 0.544 | 0.865 |
| LINE | 0.165 | **0.994** | 0.819 | 0.877 | 0.275 | 0.875 |
| DeepWalk | 0.515 | 0.971 | 0.749 | 0.923 | 0.610 | 0.906 |
| Word2Vec | 0.550 | 0.983 | 0.841 | 0.929 | 0.665 | 0.921 |
| Our Model | **0.954** | 0.892 | **0.899** | **0.950** | **0.925** | **0.923** |

According to the results on both datasets and all evaluation criteria, our model could perform better than the competing methods. This is because of using weighted PPI network and considering DC of nodes besides the network structure in learning genes’ features, employing weight-biased random walk for getting the nodes’ context, and using Relational AutoEncoder for feature embedding.

## 5. Conclusion

In detecting essential genes based on PPI network structures, extracting effective feature vectors for genes is a critical step in design of machine-learning methods. In this paper, we proposed a weight-biased random walk algorithm to get sequence of neighbors for each gene from the PPI network and generate a 4-channel feature vector for each gene from random walks-out. Then, the novel embeddings of genes were produced using a Relational AutoEncoder and were fed into an ANN classifier.

The features represented by our model gave better performance on human essential gene prediction, compared to competing methods which work only on first level of nodes’ neighborhood and analyze the network for each gene more locally. The experimental results suggest that our model is an efficient way to learn the feature representations for genes in human PPI networks.

In future works, we will focus on designing a more robust network-based method to find the feature representation of genes in the PPI network, and developing more effective machine learning methods to predict human essential genes based on these features. Additionally, since human gene can produce many different protein isoforms and there are some noises in human PPI network, improving reliability of networks would be a potential solution to improve the essential gene prediction.

## References

[1] R. Zhang and Y. Lin, “DEG 5.0, a database of essential genes in both prokaryotes and eukaryotes,” Nucleic Acids Res., vol. 37, pp. D455–D458, 2009.

[2] Y. Lu, J. Deng, J.C. Rhodes, H. Lu, and L.J. Lu, “Predicting essential genes for identifying potential drug targets in Aspergillus fumigatus,” Comput. Biol. Chem., vol. 50, pp. 29–40, 2014.

[3] G. Giaever, A.M. Chu, L. Ni, C. Connelly, L. Riles, S. Véronneau, S. Dow, A. Lucaudanila, K. Anderson, and B. André, “Functional profiling of the saccharomyces cerevisiae genome,” Nature, vol. 418, pp. 387–391, 2002.

[4] L.M. Cullen and G.M. Arndt, “Genome-wide screening for gene function using RNAi in mammalian cells,” Immunol. Cell Biol., vol. 83, p. 217, 2005.

[5] T. Roemer, B. Jiang, J. Davison, T. Ketela, K. Veillette, A. Breton, F. Tandia, A. Linteau, S. Silaots, and C. Marta, “Large-scale essential gene identification in Candida albicans and applications to antifungal drug discovery,” Mol. Microbiol., vol. 50, pp. 167–181, 2003.

[6] S. Fields and O. Song, “A novel genetic system to detect protein-protein interactions,” Nature, vol. 340, pp. 245–246, 1989.

[7] Y. Ho, A. Gruhler, A. Heilbut, G.D. Bader, L. Moore, S.L. Adams, A. Millar, P. Taylor, K. Bennett, and K. Boutilier, “Systematic identification of protein complexes in saccharomyces cerevisiae by mass spectrometry,” Nature, vol. 415, p. 180, 2002.

[8] H. Jeong, S.P. Mason, A.L. Barabási, and Z.N. Oltvai, “Lethality and centrality in protein networks,” Nature, vol. 411, pp. 41–42, 2001.

[9] G. Li, M. Li, J. Wang, J. Wu, F.X. Wu, and Y. Pan, “Predicting essential proteins based on subcellular localization, orthology and PPI networks,” BMC Bioinform., vol. 17, p. 279, 2016.

[10] B. Xu, J. Guan, Y. Wang, and Z. Wang, “Essential protein detection by random walk on weighted protein-protein interaction networks,” IEEE/ACM Trans. Comput. Biol. Bioinform., vol. 16, no. 2, pp. 377–387, 2017.

[11] W. Dai, Q. Chang, W. Peng, J. Zhong, and Y. Li, “Network Embedding the Protein–Protein Interaction Network for Human Essential Genes Identification.” Genes, vol. 11, p. 153, 2020.

[12] R.R. Vallabhajosyula, D. Chakravarti, S. Lutfeali, A. Ray, and A. Raval, “Identifying hubs in protein interaction networks,” PloS One, vol. 4, p. 5344, 2009.

[13] S. Wuchty and P.F. Stadler, “Centers of complex networks,” J. Theor. Boil., vol. 223, pp. 45–53, 2003.

[14] M.P. Joy, A. Brock, N.E. Ingber, and S. Huang, “High-betweenness proteins in the yeast protein interaction network,” J. Biomed. Biotechnol., vol. 2005, pp. 96–103, 2005.

[15] K. Stephenson and M. Zelen, “Rethinking centrality: Methods and examples,” Soc. Networks, vol. 11, pp. 1–37, 1989.

[16] P. Bonacich, “Power and centrality: A family of measures”, Am. J. Sociol., vol. 92, pp. 1170–1182, 1987.

[17] E. Estrada and J.A. Rodríguez-Velázquez, “Subgraph centrality in complex networks,” Phys. Rev. E, vol. 71, no. 056103, 2005.

[18] J. Wang, M. Li, H. Wang, and Y. Pan, “Identification of essential proteins based on edge clustering coefficient,” IEEE/ACM Trans. Comput. Biol. Bioinform., vol. 9, pp. 1070–1080, 2012.

[19] M. Li, H. Zhang, J.-X. Wang, and Y. Pan, “A new essential protein discovery method based on the integration of protein-protein interaction and gene expression data,” BMC Syst. Boil., vol. 6, no. 15, 2012.

[20] X. Tang, J. Wang, J. Zhong, and Y. Pan, “Predicting essential proteins based on weighted degree centrality,” IEEE/ACM Trans. Comput. Biol. Bioinf., vol. 11, pp. 407–418, 2014.

[21] G.T. Hart, I. Lee, and E.M. Marcotte, “A high-accuracy consensus map of yeast protein complexes reveals modular nature of gene essentiality,” BMC Bioinform., vol. 8, pp. 1–11, 2007.

[22] M. Li, Y. Lu, Z. Niu, and F.X. Wu, “United complex centrality for identification of essential proteins from PPI networks,” IEEE/ACM Trans. Comput. Biol. Bioinform., vol. 14, pp. 370–380, 2017.

[23] X.W. Tang, “Predicting essential proteins using a new method,” in Proc. International Conference on Intelligent Computing; Springer: Cham, Switzerland, 2017, pp. 301–308.

[24] W. Peng, J. Wang, Y. Cheng, Y. Lu, F.; Wu, and Y. Pan, “UDoNC: An algorithm for identifying essential proteins based on protein domains and protein-protein interaction networks,” IEEE/ACM Trans. Comput. Biol. Bioinform., vol. 12, pp. 276–288, 2015.

[25] X. Lei, S. Wang, and F. Wu, “Identification of Essential Proteins Based on Improved HITS Algorithm,” Genes, vol. 10, p. 177, 2019.

[26] Y. Chen and D. Xu, “Understanding protein dispensability through machine-learning analysis of high-throughput data,” Bioinform., vol. 21, pp. 575–581, 2005.

[27] J. Zhong, Y. Sun, W. Peng, M. Xie, J. Yang, and X. Tang, “XGBFEMF: An XGBoost-based framework for essential protein prediction,” IEEE Trans. NanoBioscience, vol. 17, pp. 243–250, 2018.

[28] F. Zhang, W. Peng, Y. Yang, W. Dai, and J. Song, “A Novel Method for Identifying Essential Genes by Fusing Dynamic Protein–Protein Interactive Networks,” Genes, vol. 10, p. 31, 2019.

[29] X. He, L. Kuang, Z. Chen, Y. Tan, and L. Wang, “Method for Identifying Essential Proteins by Key Features of Proteins in a Novel Protein-Domain Network. In Frontiers in Genetics,” Frontiers Media SA, vol. 12, 2021.

[30] C. Zhao and Z. Wang, “GOGO: An improved algorithm to measure the semantic similarity between gene ontology terms,” Scientific Reports, vol. 8, no. 1. Science and Business Media LLC, 2018.

[31] T. Blickle and L. Thiele, “A Comparison of Selection Schemes Used in Evolutionary Algorithms,” Evolutionary Computation, vol. 4, no. 4, pp. 361–394, 1996, DOI:10.1162/evco.1996.4.4.361.

[32] Q. Meng, D. Catchpoole, D. Skillicom, and P. Kennedy, “Relational Autoencoder for Feature Extraction,” in Proc. of International Joint Conference on Neural Networks (IJCNN), 2017, pp. 364–371, DOI:10.1109/IJCNN.2017.7965877.

[33] A. Odena, C. Olah, and J. Shlens, “Conditional Image Synthesis with Auxiliary Classifier GANs,” in Proc. 34th International Conference on Machine Learning Research, 2017, vol. 70, pp. 2642–2651, Available from https://proceedings.mlr.press/v70/odena17a.html.

[34] J. Wu, Q. Zhang, W. Wu, T. Pang, H. Hu, W.K.B. Chan, X. Ke, and Y. Zhang, “WDL-RF: Predicting bioactivities of ligand molecules acting with G protein-coupled receptors by combining weighted deep learning and random forest”, Bioinform., vol. 34, pp. 2271–2282, 2018.

[35] M.L. Acencio and N. Lemke, “Towards the prediction of essential genes by integration of network topology, cellular localization and biological process information,” BMC Bioinform., vol. 10, p. 290, 2009.

[36] J. Liao and K.-V. Chin, “Logistic regression for disease classification using microarray data: Model selection in a large p and small n case,” Bioinform., vol. 23, pp. 1945–1951, 2007.

[37] J. Cheng, Z. Xu, W. Wu, L. Zhao, X. Li, Y. Liu, and S. Tao, “Training set selection for the prediction of essential genes,” PloS One, vol. 9, no. e86805, 2014.

[38] K.C. Chou and H.B. Shen, “Predicting eukaryotic protein subcellular location by fusing optimized evidence-theoretic k-nearest neighbor classifiers,” J. Proteome. Res., vol. 5, pp. 1888–1897, 2006.

[39] G. Wu, X. Feng, and L. Stein, “A human functional protein interaction network and its application to cancer data analysis,” Genome Boil., vol. 11, no. R53, 2010.

[40] T. Li, R. Wernersson, R.B. Hansen, H. Horn, J. Mercer, G. Slodkowicz, C.T. Workman, O. Rigina, K. Rapacki, H.H. Stærfeldt, et al. “A scored human protein–protein interaction network to catalyze genomic interpretation,” Nat. Methods, vol. 14, pp. 61–64, 2016.

[41] F.-B. Guo, C. Dong, H.-L. Hua, S. Liu, H. Luo, H.-W. Zhang, Y.-T. Jin, and K.-Y Zhang, “Accurate prediction of human essential genes using only nucleotide composition and association information,” Bioinform., vol. 33, pp. 1758–1764, 2017.

[42] B. Perozzi, R. Al-Rfou, and S. Skiena, “DeepWalk: Online Learning of Social Representations,” in Proc. 20th ACM SIGKDD International Conference on Knowledge Discovery & Data Mining, New York, 2014, pp. 701–710.

[43] J. Tang, M. Qu, M. Wang, M. Zhang, J. Yan, and Q. Mei, “LINE: Large-scale information network embedding,” in Proc. 24th International Conference on World Wide Web, 2015, Italy, pp. 18–22.

